# Towards crystal structures of filament forming proteins

**DOI:** 10.64898/2026.02.22.707290

**Authors:** Yvette Roske, Martina Leidert, Kristina Rehbein, Anne Diehl

**Author notes:** Corresponding Author: Yvette Roske, Robert-Rössle-Str. 10, 13125 Berlin, **Email:**. **Author Contributions:** A.D., Y.R. designed research; A.D., Y.R., M.L. K.R. performed research; A.D., Y.R. analyzed data; and A.D., Y.R. wrote the paper.

## Abstract

Filament-forming proteins such as TasA (*Bacillus subtilis*) and camelysins CalY1, CalY2 (*Bacillus cereus*) pose a particular challenge for structural analysis due to their strong tendency to self-association and their polydispersity, which severely limits their ability to crystallize or to be a target for NMR-spectroscopy. To address this, it is necessary to modify the amino acid sequence to prevent filamentation. Engineering a series of N- and C-terminal truncated variants by removing flexible parts is often key to success. N-terminal extensions are also a powerful tool for obtaining crystals of fiber-forming proteins.

## Introduction

It has been shown that filament- or fiber-forming proteins are difficult to crystallize, because they strongly prefer to assemble into heterogeneous, dynamic filaments rather than single, rigid, monodisperse species, which is a prerequisite to form ordered 3D crystals. In 2018, we solved the crystal structure of TasA^1^, a filament-forming protein involved in the biofilm formation of *B. subtilis*. Several TasA constructs were required to produce the one that finally crystallized. Based on the TasA structure, we proposed models for the Camelysin CalY1 and CalY2 of *B. cereus*^1^. Recently we published our work on Camelysin of *B. cereus*^*2*^, but did not include data on our crystallization attempts. Here, we present our crystallization experiences with filament-forming proteins, such as TasA and Camelysins.

## Results

### Crystallisation of TasA

The crystallization of TasA was described in Diehl et al. 2018, but we did not clearly highlight that the construct TasA_239_ has an N-terminal extension of one single glycine besides the missing signal sequence AA 1-27 and the C-terminal AA 240-161. That protein (19.5 mg/ml) crystallizes well under the crystallization conditions of 34 % PEG 2000 MME, 0.1 M ammonium sulfate, 0.2 M lithium salicylate, 0.1 M sodium acetate pH 4.6 employing the sitting drop vapor-diffusion method, examples of resulting crystals are shown below (Fig.1).

**Figure 1.**
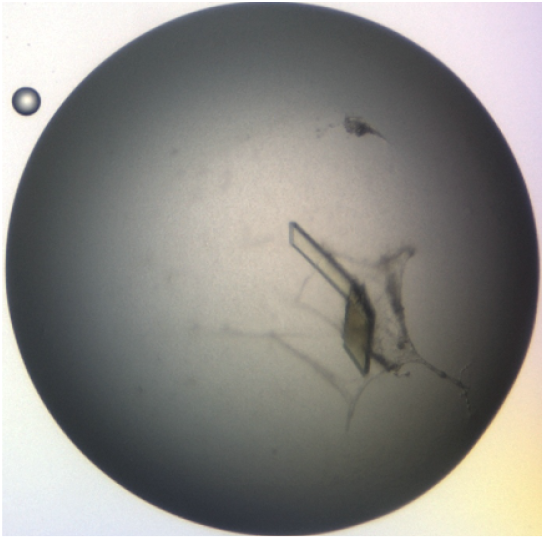
Crystals of TasA_239_ grown at 20 °C within 2-6 days diffracting to 1.8 Å.

### Crystallization attempts of CalY1 and CalY2

Crystallization attempts were performed with CalY1 and CalY2 in 20 mM Tris HCl buffer pH 7.0 with 150 mM NaCl using the sitting-drop vapor-diffusion method. A Gryphon pipetting robot (Matrix Technologies Co.) was used to pipette 200 nL of protein at a concentration of 17 to 26 mg/mL into an equal volume of precipitant solution. Commercially available crystallization screens with total ∼700 conditions were applied. The Rock Imager 1000 storage system (Formulatrix) was used for storing and imaging of the experiments at 20 °C. Details of the used camelysin variants and concentrations are summarized in Table 1. The observed strong tendency of mature CalY2 to become gel-like was reduced by N-terminal extensions with a single glycin (G) or serin and alanine (SA) or truncations starting from AA 42, allowing crystallization experiments to be performed. Needles were obtained for CalY1_42-193 in 25 % PEG 3350, 0.1 M sodium citrate pH 3.5 (blue in Table 1) and for CalY2_G_28-195 in 5-11 % 2-Propanol, 0.1 M MES pH 6 – pH 7 (green in Table 1) (Fig.2A, B). Polydispersity of CalY1 and its interconversion into different species prevents crystallization^2^. Microcrystals were found for the N-terminal extended (SA) and C-terminal truncated CalY2_SA_28-189 using 5 %-11 PEG 4000 or PEG 8000 und 0.1 M Bicine pH 8.5 (red in Table 1) (Fig. 2C) Unfortunately, optimization of the crystallization conditions did not lead to proper diffracting crystals suitable for structure determination.

**Table 1:** Overview of all CalY1 and CalY2 variants checked for crystallisation.

|  | N-terminale<br>extention | AA | C-terminale<br>extension | Concentration<br>mg/ml | vector | cleavage |
| --- | --- | --- | --- | --- | --- | --- |
| CalY1 II |  | 29-197 |  | 19.8 | pET-HisSumo_TEV_EK<br>CalY1_29-197 | EK |
| CalY1 II | SADDDK | 29-197 |  | 22 | pET-HisSumo_TEV_EK<br>CalY1_29-197 | TEV |
| CalY1 |  | 42-193 |  | 23 | pET-HisSumo_CalY1_42-193 | Sumo |
| CalY2 | SA | 28-195 |  | 17,4 | pET-HisSumo_TEV_<br>CalY2_28-195 | TEV |
| CalY2 | G | 28-195 |  | 16 | pET-HisSumo_TEV(G)_<br>CalY2_28-195 | TEV |
| CalY2 | G | 28-195 |  | 15 | pET-HisSumo_TEV(G)_<br>CalY2_28-195 | TEV |
| CalY2 |  | 28-195 | GFP | 16,6 | pET-HisSumo_<br>CalY2_28-195_GFP | Sumo |
| CalY2 | G | 28-189 |  | 11,7 | pET-HisSumo_TEV(G)_<br>CalY2_28-189 | TEV |
| CalY2 | G | 28-189 |  | 11,3 | pET-HisSumo_TEV(G)_<br>CalY2_28-189 | TEV |
| CalY2 | SA | 28-189 |  | 12 | pET-HisSumo_TEV_<br>CalY2_28-189 | TEV |
| CalY2 |  | 42-189 |  | 10,7 | pET-HisSumo_<br>CalY2_42-189 | Sumo |
| CalY2 |  | 42-189 |  | 17,7 | pET-HisSumo_<br>CalY2_42-189 | Sumo |

**Figure 2.**
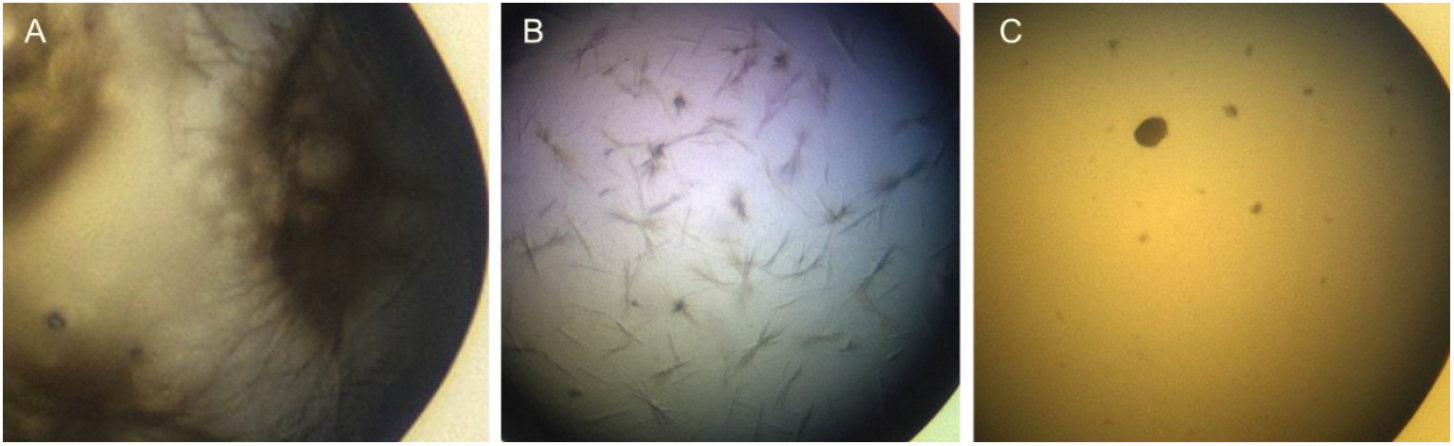
Crystals of Camelysin constructs. **A)** Thin needle crystals could be achieved of CalY1_42-193 within 1-3 days. **B)** Needle bundles of CalY2_G_28-195 appeared within 10 days. **C)** CalY2_SA_28-189 yielded microcrystals between 3 to 7 days.

## Discussion and Conclusion

In addition to N- and C-terminal truncations, N-terminal extension can also promote the crystallization of filament-forming proteins. For TasA, just one amino acid was enough to prevent filamentation and produced well-diffracting crystals. However, to stop the rapid filamentation of CalY2, an N-terminal extension containing SA or a single glycin residue (G) was required to produce microcrystals or small needle bundles, respectively. CalY1_42-193 showed fuzzy needle-like structures out of precipitated protein and was very difficult to optimize. In our hands, CalY2_G_28-195 proved to be the most promising candidate with regard to crystallization. Variations in the crystallization conditions did show some changes in the crystallization behavior, but not to an extent that would have allowed us to produce individual diffracting crystals. Crystallization at different temperature may improve the results. As no further work is planned in this regard from our site, we would like to share these unpublished results here with the interested community, as these modification strategies demonstrate that terminal flexibility and native polymerization interfaces can be decoupled by minimal sequence changes. This enables the structural analysis of otherwise filament-forming proteins as shown for TasA.

## Material and Methods

### Constructs

Basic constructs for TasA have been described elsewhere^1^. Starting from pCA528_His_Sumo_TasA28-261 we generated a vector for TasA_239_ production by mutation of K240 (AAG) to a stop codon TAG. Additional we introduced the coding region for a TEV cleavage site by excluding SUMO. So that on protein level after TEV-cleavage a Glycin remains at the N-terminus of TasA_239_.

For CalY1 (197 AA, Q8GJ76; Q8GJ76_BACCE; AF-Q8GJ76-F1) and CalY2 (195 AA, A0A0E3SV09_BACCE; AF-A0A0E3SV09-F1) codon-optimized strings were synthesized by GeneArt (Regensburg, Germany). The 5’and 3’ends flanking the open reading frame contain BsaI sites to generate pET-based expression vectors carrying a kanamycin resistance cassette and a T7 promotor via an in-house Modular Cloning System (provided by Martin Bommer / MDC). We came up with 2 vectors coding for HisSumo_TEV_EK_CalY1_29-197_3C_GFP (AA F28 was accidently missing), and HisSumo_TEV_CalY2_28-195_GFP. All variants for TasA, CalY1 and CalY2 were done by modified QuikChange site directed mutageneses. Table 2 shows all used primer synthesized by BioTez Berlin/Germany. Resulting constructs were confirmed by sequencing service at LGC genomics Berlin/Germany.

**Table 2:**
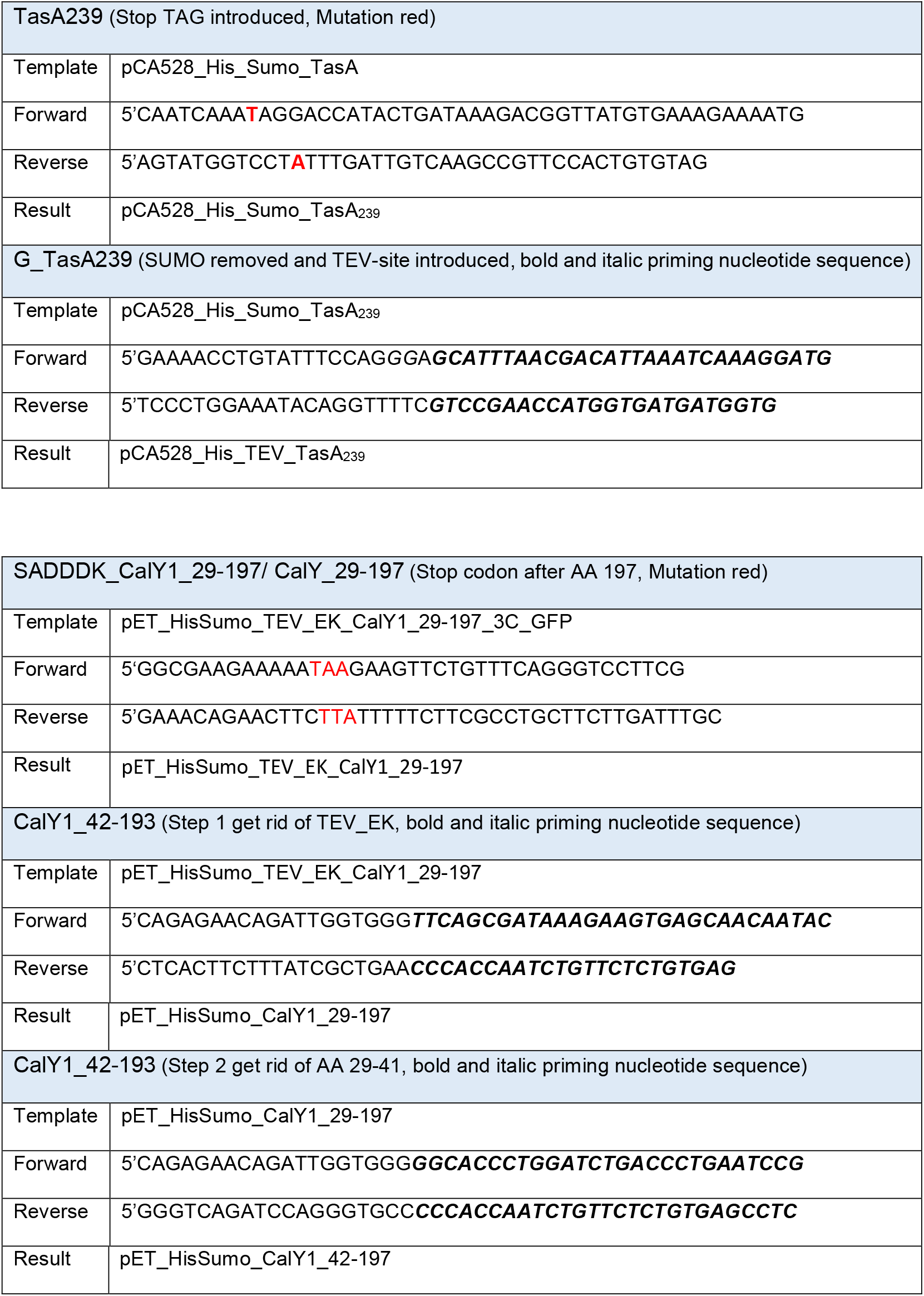
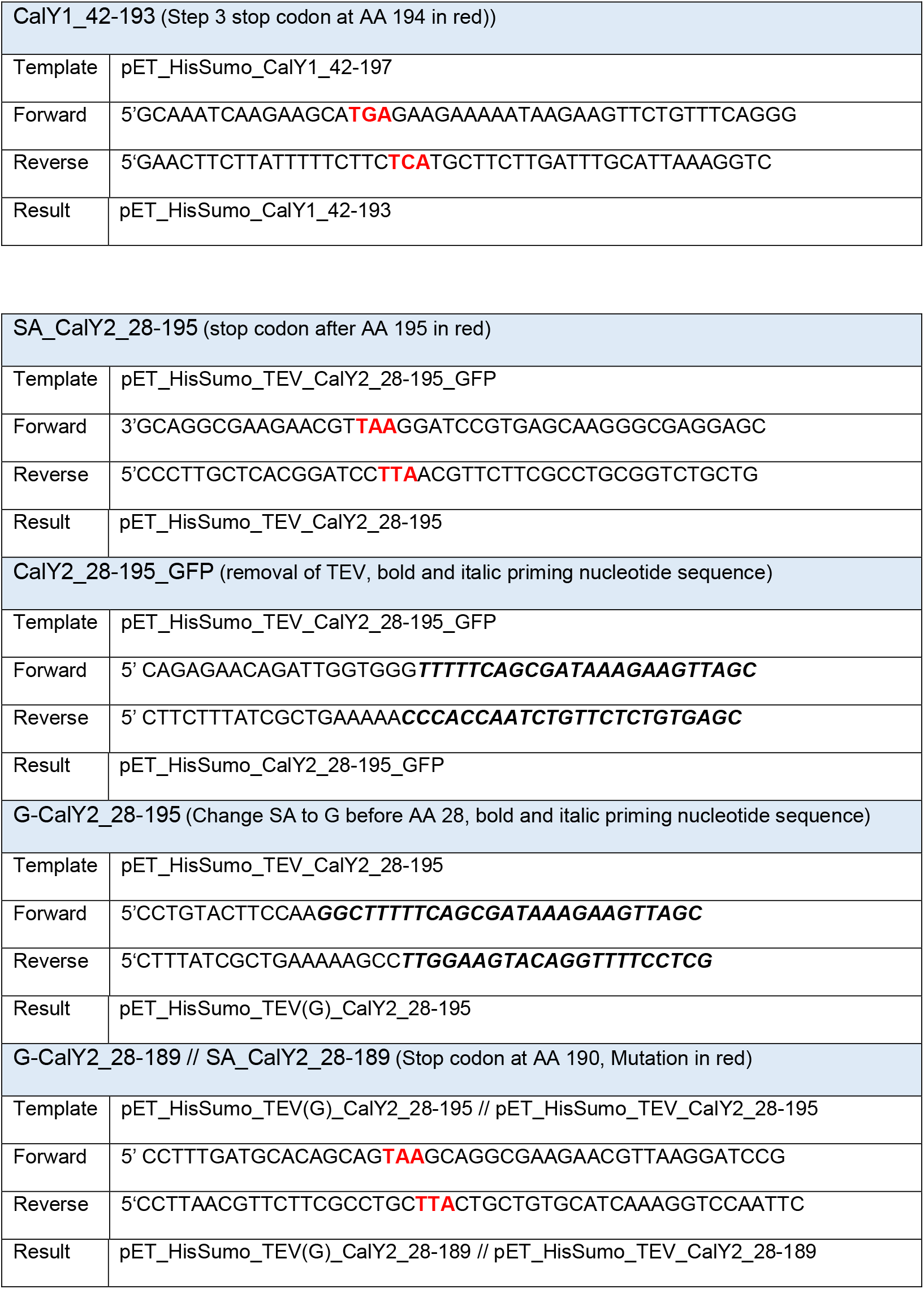
Primer used for vector creation to produce TasA_G28-239 and the proteins in Table 1.

### Recombinant production

Protocols for TasA were described before^1,3^. HisSumo-Fusions CalY1 and CalY2 yield well-soluble protein after expression on rich and minimal medium. Approximately 60 - 80 mg of fusion protein were obtained from a 1 L culture after the first metal chelate (MC) column. Following cleavage of the HisSumo tag vector dependent by Sumoprotease, TEV or EK under dialysis, a second MC column was used to remove the Fusionpartner. If possible the CalYs were concentrated and applied to a 120 or 320 ml Superdex 75 gel filtration column^2^.

### Crystallization

All crystallization attempts for the Camelysin variants were done at 20 °C with a drop volume of 200 nl protein solution and equal volume precipitant solution in a 96-well crystallization plate using the sitting drop vapour-diffusion method. The droplets were images at day 0, 1, 3, 7, 14, 21, 28, 48.

## Abbreviations

AA: amino acid(s)
CalY: Camelysin
PEG: Polyethylene glycol
MME: monomethyl ether
MES: 2-(N-morpholino)ethanesulfonic acid, EK, enterokinase cleavage site
TEV: Tobacco Etch Virus cleavage site
3C: PreScission protease cleavage site
GFP: Green Fluorescent Protein

